# Scales of marine endemism in oceanic islands and the Provincial-Island endemism

**DOI:** 10.1101/2024.07.12.603346

**Authors:** Hudson T. Pinheiro, Luiz A. Rocha, Juan P. Quimbayo

## Abstract

Oceanic islands are remote environments commonly harboring endemic species, which often are unique lineages originated and maintained by a variety of ecological, biogeographical and evolutionary processes. Endemic species are found mostly in a single oceanic island or archipelago, however, a great number of species can be considered multiple-island endemics, i.e. species found on multiple oceanic islands that still have a restricted distribution. The geographic criteria chosen to classify endemic species has a direct impact on the endemism rate of islands, and many studies have used multiple scales of endemism (single and multiple-island endemics), which has historically influenced wide-scale comparisons. In this perspective, we accessed the importance of single and multiple-island endemic species to the biodiversity of oceanic islands, introducing the concept of Provincial-island endemism as an additional strategy to standardize biogeography studies.

## Main Text

It is well known that many oceanic islands harbor unique biodiversity, and due to their faunal simplicity and limited distributions, these endemic species have often been the focus of important evolutionary, ecological and biogeographic studies. The diversification patterns and premises in island biogeography, however, differ for terrestrial and marine realms. Terrestrial species often diversify following adaptive radiation, while marine species, which are better dispersers, experience fewer opportunities for in situ cladogenesis (Pinheiro et al., 2017). Therefore, marine endemism is more often related to geographical isolation (Ávila et al., 2019, 2018; Hachich et al., 2019, 2015). Accordingly, the origin of new lineages based on the colonization of islands by immigrants and their subsequent isolation represents one of the most classical scenarios of evolution, a process known as peripatric speciation. This process has been suggested to drive the highest levels of endemism in islands of both Pacific (e.g., Hawaii and Eastern Island) and Atlantic (e.g., Ascension, St Helena and St Paul’s) oceans (Delrieu-Trottin et al., 2019; Floeter et al., 2008; Hourigan and Reese, 1987; Randall, 1998). The colonization process is suggested to happen stochastically, but also in waves, what is often related to favorable oceanographic conditions for founder populations to reach the islands or source populations to expand (Delrieu-Trottin et al., 2017; Hodge et al., 2014; Pinheiro et al., 2017).

Recent studies have explored a variety of processes that contribute to the current endemism level of oceanic islands. Ecological or parapatric speciation has been evoked to explain the high endemism rate in the Marquesas (Gaither et al., 2015; Randall, 1998), and the evolution of species in the Northeastern Brazilian oceanic islands (Rocha et al., 2005). Pleistocene sea-level variations have also promoted vicariant events, driving allopatric speciation (Pinheiro et al., 2017) – the connections between source and insular populations change as shallow habitats are created or suppressed. In the Vitoria-Trindade Chain, low sea level stands exposed many seamounts that function as stepping-stones for weak disperser species to reach remote oceanic islands (Simon et al., 2022). Subsequent sea-level rise isolated their populations eliminating connecting shallow habitat, closing migration and driving allopatric speciation. Moreover, phylogenetic studies are also revealing many paleoendemic species in oceanic islands (relict lineages that likely had wider distributions in the past; Delrieu-Trottin et al., 2019; Pinheiro et al., 2017).

In this context, islands function as dynamic biodiversity engines that not only work as cradles for species origination, but also as museums that contribute to lineages persistence over time. The role of islands to develop and support new lineages is commonly measured by their endemism rate, which is the proportion of the local biodiversity that is exclusively found there. Recent empirical work and biogeographic analyses have shown that many islands share species exclusively found in remote and isolated oceanic islands (Floeter et al., 2008; Pinheiro et al., 2018; Robertson and Cramer, 2009). These multiple-island endemic species have their origin or persistence associated with the insular environment, and in many cases, they can compose a substantial portion of the local biodiversity (Pinheiro et al., 2020) and characterize provinces (Robertson and Cramer, 2009). In this perspective we assess the importance of single and multiple-island endemic species to the biodiversity of oceanic islands, discuss how the scale of endemism has historically biased large-scale biogeographic comparisons, and suggest strategies to move forward. We are herein considering as oceanic islands only remote formations predominantly formed by volcanic activity, and for simplification, we will hereafter refer to oceanic islands, archipelagoes, atolls, and shallow seamounts, all as “oceanic islands”.

We analyzed the distribution of 7,289 fish species associated with reef environments of 87 oceanic islands and 189 coastal reefs around the world (e.g., Kulbicki et al., 2013), and uncovered that 5,260 (72%) species occur in oceanic islands. According to their composition, the islands are strongly structured by ocean basins, where islands of the Atlantic Ocean are more similar among themselves, while a large gradient of similarity is observed in the Indo-Pacific (Figure 1). This gradient in similarity levels seems to be driven mainly by a combination of the islands’ isolation and distance from the Coral Triangle, which result in two compositional patterns (Randall, 1998): First, a gradient in species richness and associated nestedness occurs, with remote locations exhibiting a strong selective pressure favoring species with superior dispersal capabilities. Second, higher levels of endemism are observed in these remote locations, resulting from speciation in isolated populations. Further, scrutinizing the composition of the islands we observed that 4,346 species occur in both oceanic islands and continental coastal reefs, and 890 species are exclusively found around oceanic islands. Thus, 12.2% of the world’s reef fish biodiversity seems to be endemic to oceanic islands, where 541 (60.7%) are single island endemics, while 349 (39.3%) are multiple island endemics (Figure 2A). Therefore, both widespread and island restricted species contribute to the similarity among locations (Figure 1).

**Figure 1.**
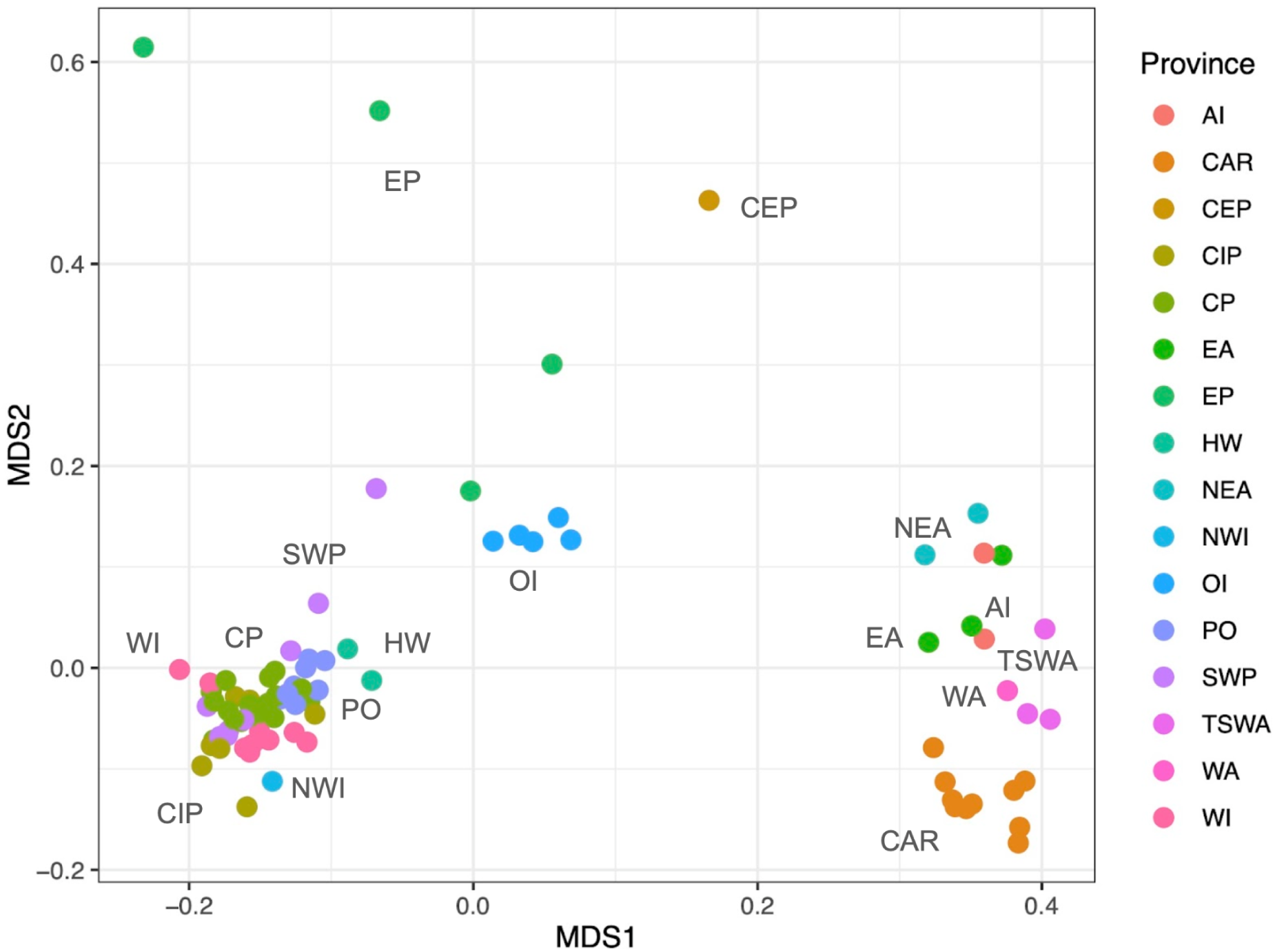
Similarity among the oceanic islands analyzed, based on the Jaccard coefficient and presence-absence data of the distribution of 5260 reef fish species. The biogeographic provinces used in this analysis were adapted from The provinces presented were adapted from Kulbicki et al. (2013): AI - Atlantic Islands; CAR - Caribbean; CEP - Continental TEP; CIP - Central Indo-Pacific; CP - Central Pacific; EA - Eastern Atlantic; EP - Eastern Pacific; HW - Hawaiian; NEA - North-Eastern Atlantic; NWI - North-Western Indian; ETP - Oceanic Islands Eastern Tropical Pacific Pacific; PO - Polynesian; SWP - South-Western Pacific; TSWA - Tropical South-Western Atlantic; WA - Western Atlantic; WI - Western Indian Ocean.

**Figure 2.**
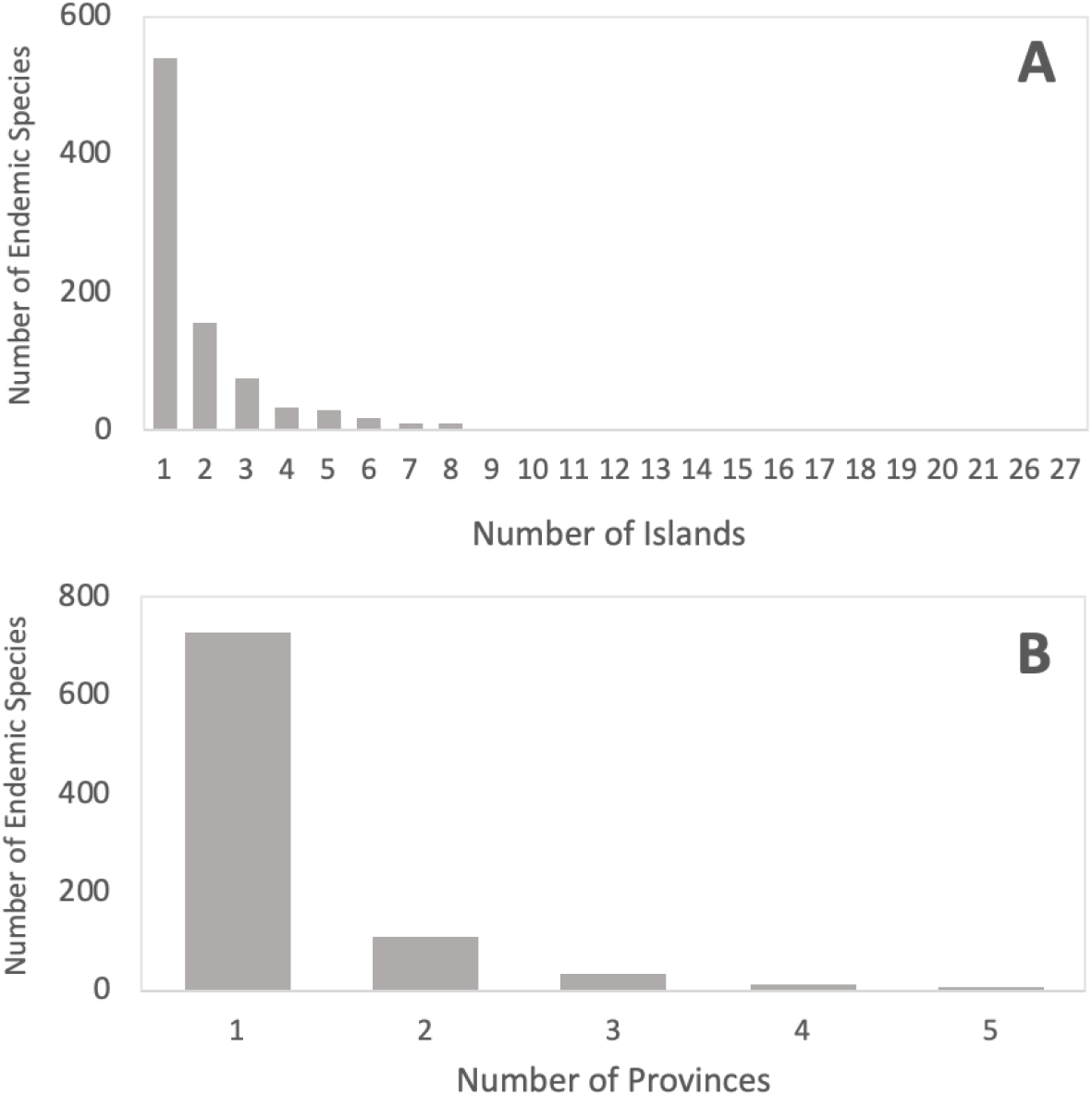
Distribution of endemic species among islands and provinces. A) Relationship between the number of island endemic species and the number of islands they occur. B) Relationship between the number of island endemic fishes and the number of provinces they occur in.

Regarding multiple island endemics, some species are genetically connected in distances that are greater than their isolation to the continent (Rocha et al., 2005; Bernardi et al., 2014). For instance, *Stegastes beebei* is an endemic species found in Galápagos, Cocos, and Malpelo, in the Eastern Tropical Pacific, but not in the adjacent coastline (Robertson and Cramer, 2009). Similarly, the oceanic islands in the Mid-Atlantic Ridge, over 1,000 km apart, also share many endemic species (Brown et al., 2019; Pinheiro et al., 2020; Wirtz et al., 2017). In the Brazilian Province, there is an extreme case where a species (*Opistognathus thionyi*) is shared between islands almost 2,000 km apart, where the shortest distance to the mainland is only 350 km (Pinheiro et al., 2018) – a connectivity pattern also observed for a coral lineage (Peluso et al., 2018) and a Brazil-Caribbean link (Rocha et al. 2005). It has been suggested that an oceanic “pathway” driven by stochastic and ecological factors connects species among oceanic islands at the same time that constrains their establishment in the continental shelf (Pinheiro et al., 2020, 2018). Indeed, several species might have the ability to disperse, but the environment is a key factor for their persistence (Rocha et al., 2005; Mazzei et al., 2021), and can contribute for the establishment of historical refuges in oceanic islands during unfavorable geological periods (Peluso et al., 2018). Thus, even the most remote archipelagoes do not constitute dead ends, often exporting new lineages to islands and seamounts nearby (Simon et al., 2022) and also back to biodiversity centers in a process known as Biodiversity Feedback (Bowen et al., 2013).

From these multiple-island endemic species, 231 (66%) are found in only two or three islands or archipelagoes, and a strong negative relationship is observed between the number of endemic species and the number of islands they occur (Figure 2). For instance, very few multiple-island endemic fishes are found in more than 10 islands, so, in general, they tend to have restricted distributions. This pattern can be explained by the evolutionary history and life-history of endemic fishes, as well as the geography of islands. Fishes in general have good dispersal potential due the combination of pelagic larval duration and swimming abilities, and most species that inhabit oceanic islands or are widespread along the oceans have large body size, pelagic spawning and larvae, and exhibit rafting abilities (Luiz et al., 2013, 2012; Mazzei et al., 2021; Pinheiro et al., 2018). The evolution of an endemic lineage often occurs after the termination of gene flow with source populations, a process commonly observed in vicariant events and stochastic colonization of weak dispersers, or the establishment of weak and moderate dispersers in extremely remote locations. This combination of life-history traits and island geography contribute to a wide variety of morphologies and taxonomic groups among endemic reef fishes (Robertson, 2001), and to the positive relationship between the number of endemic fishes and the island isolation (Hachich et al., 2015). Therefore, the same drivers that allow speciation can also contribute to the maintenance of their restricted distribution. Moreover, many lineages are considered neo-endemics, evolving during the Pleistocene and possibly not having enough time to disperse or adapt to other oceanic islands and seamounts. For instance, the evolutionary history of endemic fishes in Rapa Nui (Pacific Ocean) and Trindade (Atlantic Ocean) islands shows a majority of neo-endemics with restricted geographic range, while some wider-distributed endemics show older lineages (Delrieu-Trottin et al., 2019; Pinheiro et al., 2017).

The distribution of endemic fishes contributes directly to the endemism rate of the islands. The Hawaiian archipelago and Rapa Nui have traditionally been considered hotspots of endemism, showing endemism rates above 20% (Randall, 1998). However, according to our analyses, these localities would present 10 and 13.4% of endemism, respectively, if only single-island endemics are considered (Figure 3). The endemism rates of these locations are substantially higher when multiple-island endemics are considered, rising to 22.2 and 35.9% respectively. Some islands that present high proportions of multiple-island endemics do not necessarily present high proportions of single-island endemics. Indeed, most of the islands herein analyzed (66.6%) present more multiple-island than single-island endemics, and 25.4 % of the islands do not have a single-island endemic fish, some presenting high proportions of multiple-island endemics, such as Salas y Gomes (32.0%), Johnston Atoll (16.2%), Madeira (7.1%) and Gambier (7.0%). Accordingly, considering endemic species that are shared among nearby islands, the highest endemism rates are found in the remote islands of Juan Fernandez and Desventuradas, where 69.5 and 53.3% of their respective biodiversity are composed by island endemics restricted to their biogeographic province. In the Atlantic, over 24% of the fishes found in each of the islands of St Helena and Ascension are exclusively found in the Mid-Atlantic Ridge.

**Figure 3.**
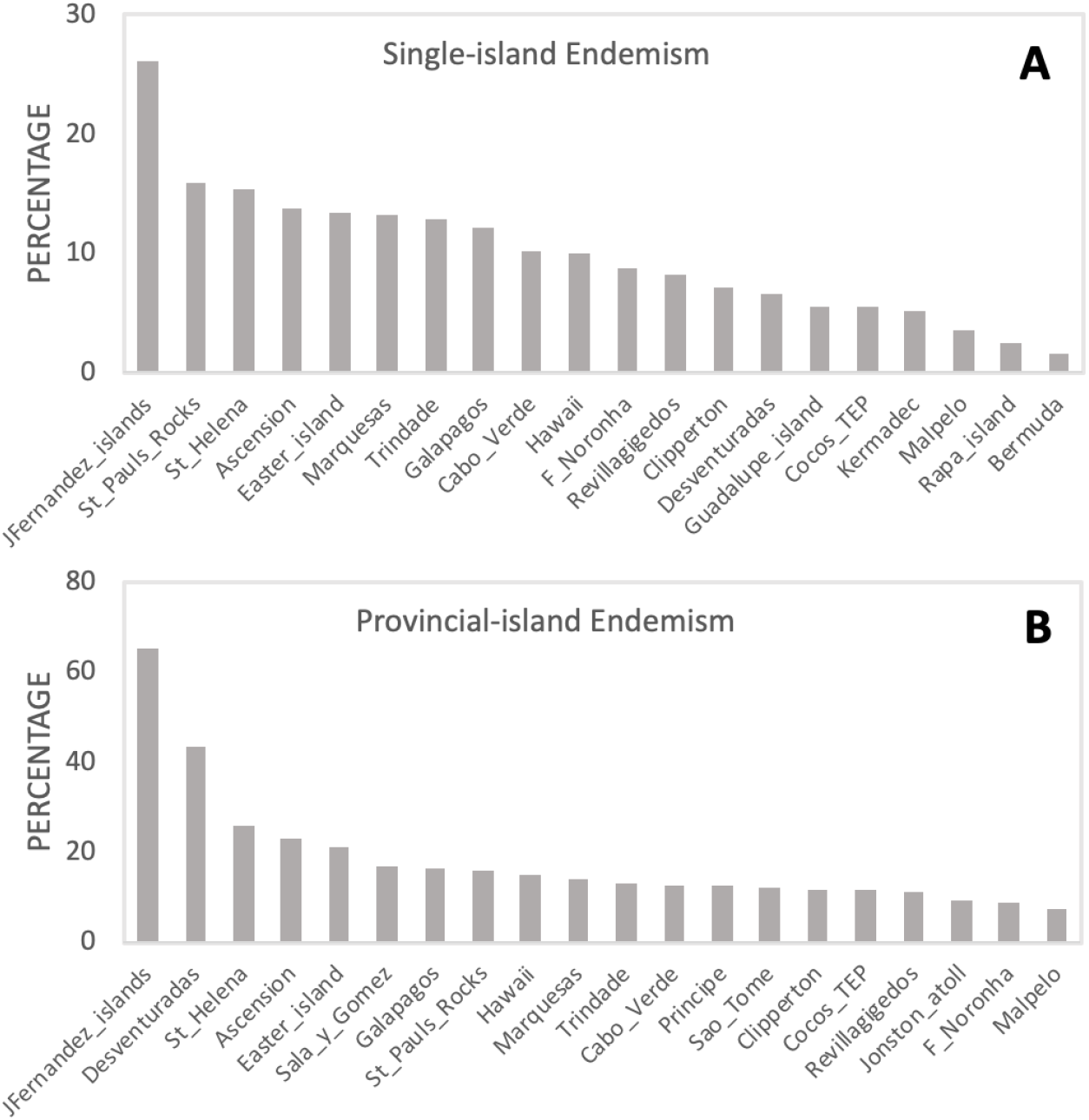
Proportion of single-island (a) and province-island (b) endemism among the oceanic islands with highest endemism.

So, why do the scales of endemism in oceanic islands matter? Some of the reasons include their importance for evolutionary, ecologic and biogeographic studies, as well as for conservation planning. Because they contain simplified species communities, oceanic islands have for a long time been considered natural laboratories to understand the origin of marine biodiversity, and omitting about 40% of their endemicity represent disregarding a great amount of evolutionary history and diversification processes that are promoted in the islands. Recent studies that have included multiple island endemics in their analyses are finding a wide range of genetic connectivity, meaning that, while some are expanding their distribution, others are under ongoing fragmentation (Bernardi et al., 2014; Delrieu-Trottin et al., 2017). Other studies are revealing relict species, remains of apparently previously widely distributed lineages that only survived in remote locations. Moreover, the definition of some subprovinces, such as the Ocean Island Province in the Tropical Eastern Pacific, was partly based on single and multiple island endemism (Robertson and Cramer, 2009). Therefore, the inclusion of single and multiple island endemics in evolutionary and biogeography studies is key to clarify our understanding of the evolution of marine communities in remote islands.

All oceanic islands analyzed showed some degree of endemism (i.e., containing single or multiple island endemic species), having many species that are exclusively adapted to oceanic environments. The adaptation process that leads to speciation or distributional expansion is also associated with ecological advantages. These species might be better competitors and have higher abundances than widespread species (Hobbs et al., 2012; Macieira et al., 2015). The ecology of the endemic fishes can contribute insights about their dispersive ability (Robertson, 2001) and be related to their evolutionary history (Pinheiro et al., 2017). Thus, considering the endemism status of a species could have important implications on ecological studies that use these remote environments as natural laboratories.

In biogeography, overlooking endemics that are shared between islands can downplay the strength of barriers and/or the ecological factors that drive speciation and taxonomic uniqueness. Endemism represents singularity, a proportion of the fauna that evolved or has been maintained in a certain province, region or location. Notwithstanding, studies have traditionally been reporting endemism rates for islands using multiple scales of endemism (single and multiple-island endemics), which has historically biased wide-scale contrasting. To move forward in the comparisons among oceanic islands, we suggest biogeographers to clarify two aspects of endemism: 1) single-island endemics; endemism rate of the local biodiversity when removing species recorded in different oceanic islands; 2) provincial-island endemics; local endemism rate considering as endemic those species restricted to oceanic islands of the same biogeographic province (e.g., provinces in Kulbicki et al., 2013). Following the former criteria, the islands with highest endemism rates are Juan Fernandez, St. Paul Rocks, St. Helena, Ascension, Eastern Island, Marquesas, Trindade, Galápagos, Cabo Verde and Hawaii, all with over 10% of single-island endemism (Figure 3A). According to the proposed novel criteria, the islands of Juan Fernandez, Deventuradas, St Helena, Ascension, Rapa Nui, Salas y Gomes, Galápagos, St. Paul Rocks, Hawaii, and Marquesas present the highest endemism rates, ranging from 18.5% to 69.5, (Figure 3B). Applying these criteria leads to better assessments of the drivers of speciation and species maintenance.

Although reef fish communities in oceanic islands have experimented severe local impacts by both commercial and recreational fisheries (Guabiroba et al., 2020; Luiz and Edwards, 2011; Pinheiro et al., 2010; Quimbayo et al., 2019; Tuya et al., 2006), very few endemic species are considered important fishing resources.. Endemic lineages of angelfishes of unique color morphs from St. Paul’s Archipelago have reached the aquarium trade, but they are now protected. Nevertheless, the restricted distribution makes endemic fishes vulnerable to population fluctuations and crashes, and global stressors driven by climate change, ocean warming and acidification are ongoing issues. For instance, *Azurina eupalama*, an endemic species from Galápagos Archipelago, hasn’t been seen since the 1982/83 very strong El Nino (Edgar et al., 2010). Many endemic species show signs of bottlenecks and decreasing population size (Pinheiro et al., 2017), which could make them more vulnerable to human derived impacts. The IUCN threat classification has historically used the distribution of species and their population sizes as a proxy for vulnerability, and a special attention to single and multiple-island endemics is here supported.

Moreover, biodiversity hotspots have been determined based on the endemism rate and threats to their biodiversity, and have often been used as a proxy for the establishment of priority areas for conservation (Roberts et al., 2002). This view has evolved to the protection of both connectivity and genetic diversity as well as peripheral areas of high endemism (Hughes et al., 2002). Many very-large marine protected areas have been created around oceanic islands throughout the world, however, many of them do not directly protect coastal reefs and endemic species, contributing to the mismatch between protection and the conservation priorities (Giglio et al., 2018; Lindegren et al., 2018). Therefore, understanding how much of the diversity of an island represents species of restricted distribution is the first step to clarify its vulnerability and priority for conservation. Applying current scientific criteria in conservation planning for the protection of priority areas is the next step to be followed.

## Materials and Methods

The data used in this study was extracted from Quimbayo et al. (2022). To quantify biogeographical organization, we assessed species composition similarity among islands. Specifically, we estimated the Jaccard coefficient using presence-absence data for 5,260 reef fish species. This estimation was performed with the “dist” function from R package *vegan* (Oksanen et al., 2020). We then summarized the similarity matrix using a Non-Metric Multidimensional Analysis (nMDS) implemented with the “metaMDS” function from R package *vegan* (Oksanen et al., 2020).

## Acknowledgements

We thank the administrative and logistics support of CEBIMar-USP, CalAcademy and UM and their staff. We are grateful for the support of donors who endorsed the California Academy of Sciences’ Hope for Reefs initiative, Rolex Award for Enterprise, Fundação de Amparo à Pesquisa do Estado de São Paulo and Fundação Grupo O Boticário de Proteção à Natureza for funding expeditions throughout oceanic islands of the Pacific, Indian and Atlantic oceans.

## Data, scripts, code, and supplementary information availability

Data is available at Quimbayo et al. (2022) A trait-based approach to marine island biogeography. https://zenodo.org/records/7316869.

